# Single-Molecule FRET and Tracking of Transfected Biomolecules in Living Cells

**DOI:** 10.1101/2023.09.15.557875

**Authors:** Abhinaya Anandamurugan, Antonia Eidloth, Veronika Frank, Philipp Wortmann, Lukas Schrangl, Chenyang Lan, Gerhard J. Schütz, Thorsten Hugel

## Abstract

Proteins and DNA in cells exhibit different conformational states, which are influenced by dynamic interactions with other biomolecules. All these interactions are affected by the molecules’ localization within the cell, i.e., their compartmentalization. Such, in cellula, compartment-specific dynamics is difficult to measure, because of limitations in instrumentation, autofluorescence of cells, and the necessity to track diffusing molecules. Here, we present a bottom-up engineering approach, which allows us to track transfected proteins *in cellula* and to analyze time-resolved single-molecule FRET efficiencies. This has been achieved by alternating laser excitation (ALEX) based three-channel (donor, acceptor and FRET) tracking with a live-cell HILO microscope. We validate our strategy by characterizing long-term static-FRET traces of customized DNA with known dye positions. We utilize two different transfection strategies, namely a biological (Streptolysin-O toxin protein) and a physical one (Microinjection). By comparing in vitro and in cellula measurements we show that the cellular environment in this case changes the FRET efficiency by about 25%. In addition, we evaluate single-molecule FRET traces for the heat shock protein Hsp90 in cellula. The obtained FRET efficiency distribution is largely consistent with known Hsp90 structures and *in vitro* distributions, but also shows some clear differences. Altogether, we show that FRET-TTB (Förster Resonance Energy Transfer-Tracking of Transfected Biomolecules) opens the path to study protein state changes of transfected biomolecules in cellula, including time-resolved cellular localization.

**Significance:** Inside cells, proteins and DNA can change shape depending on their environment and their interactions. Studying these changes for single biomolecules is challenging because they move around in the cell and the cell environment makes it hard to see them clearly. Here, we demonstrate a new technique called FRET-TTB that enables us to track individual proteins inside living cells over time using single-molecule fluorescence microscopy. We validate the approach with custom DNA and apply it to study the heat shock protein Hsp90. Our results show that this method can reveal changes in protein conformation and location within living cells.

## Introduction

Investigation of protein structure and dynamics *in vitro* has largely enhanced our understanding of proteins and their interactions. Determining how protein structure, dynamics, and interactions are affected in a crowded non-equilibrium environment, like the cytosol of living cells, in a spatially resolved way poses the next challenge. *In vitro*, long-term smFRET measurements have been most reliable in detecting dynamic distance changes of less than 10 nm [1]. Consequently, a promising direction is to extend smFRET measurements to live-cell applications. For some membrane proteins, this has already been achieved successfully [2], but not for the recruitment and release of proteins to and from membranes [3] or for cytosolic proteins like for example Hsp90. The central problem to achieve this in mammalian cells is to track single molecules showing conformational dynamics and to disentangle the effect of the microenvironment from protein conformational state changes. Two main types of smFRET methods have been employed to study conformational changes: lifetime photon burst measurements with confocal microscopy, and fluorescence intensity measurements with total internal reflection (TIR) microscopy or Highly Inclined Laminated Optical Sheet (HILO) microscopy. Lifetime measurements provide rich information on molecular decay times and conformations [4], but they are not suitable for long-term measurements following one single molecule over time, especially if this diffuses in the cell. Standard two-color smFRET is generally suitable for long-term measurements on a single molecule, but only if the molecule is immobilized [5]. Therefore, it can also not be directly applied for mobile molecules in living cells. Fluorescence Correlation Spectroscopy (FCS) on the other hand has been widely used to study diffusion in biological systems [6], but is application to study conformational changes is still in its infancy.

The challenge is to localize both single fluorophores, the donor dye and the acceptor dye, track them and determine the FRET efficiency and stoichiometry over time. Here we present a method that meets this challenge *in cellula* by establishing long-term smFRET measurements of the cytosolic protein Hsp90 using HILO microscopy [7], three-channel tracking (Donor, FRET, Acceptor)[8] and cell transfection. The three channels are the result of alternating laser excitation (ALEX) [9], [10], which is necessary to recognize the fluorophores’ photophysics. Here and in the following, the Donor signal is the signal intensity in the donor channel upon donor excitation (*I*_Dem|Dex_), FRET the intensity in the acceptor channel upon donor excitation (*I*_Aem|Dex_) and Acceptor the intensity in the acceptor channel upon acceptor excitation (*I*_Aem|Aex_) Key requirements for tracking and obtaining long-term intensity traces of single molecules are photophysically stable FRET dye-pairs and low intracellular concentrations of the labeled molecules. This enables clear separation of signals, detectable for a long period of time. Our strategy to achieve this is *in vitro* labeling of purified proteins with synthetic high-performance dyes and the subsequent transfection of cells with the pre-labeled molecules. This allows for a high degree of control over the final amount of labeled molecules inside the cell and therefore helps to maintain single-molecule concentrations [11]–[13]. Additionally, and in contrast to endogenously expressed fluorescent proteins, the cell’s native cellular expression is preserved. We tested and compared two mechanistically different transfection strategies: a biological method using the pore-forming toxin Streptolysin-O (SLO)[14]–[17] and a physical method, namely microinjection [4], [18], [19]. Both have the capability to introduce biomolecules with a high molecular weight like DNA or the yeast Hsp90 homodimer with a molecular weight of about 82 kDa per monomer. We first benchmark the power of our FRET-TTB approach and the transfection strategies by quantifying single-molecule FRET of customized DNA oligos with known dye distances. We then show that this approach with SLO transfection also works for the molecular chaperone and heat shock protein Hsp90. Due to its role as a molecular chaperone, the resident cytosolic Hsp90 presents a hub for protein interactions and is involved in multiple proteomic pathways [20], [21]. Notably, *in vitro* single-molecule studies showed that the Hsp90 dimer occupies two major, dynamically exchanging conformational states, namely an N-terminal open and a closed state [22], [23]. However, the relation between the dynamic conformational state changes and Hsp90s biological functions is still elusive [24]. Live-cell smFRET measurements are ideal for addressing this, enabling to watch Hsp90 at work in its native environment. Employing our approach we were already able to follow single molecules inside living HeLa cells for up to 89 seconds and quantify smFRET for more than 20 seconds. A comparison of the quantified live-cell FRET efficiencies with *in vitro* results and theoretical FRET efficiencies shows good agreement and therefore proves the method successful. This opens up new ways to study how proteins function in their natural environment.

## Materials and Methods

### DNA and Protein Preparation

To understand the quality of the obtained smFRET traces, we first analyzed standard DNA, previously used in *in vitro* smFRET experiments and benchmarked in [25]. To prevent intracellular degradation, we used modified, end-sealed DNA, labeled with Lumidyne dyes LD555 and LD655 (*ATD Bio*). Here, the ends of the DNA oligo are ligated with the use of copper-based click-chemistry. Yeast Hsp90 was designed with a C-terminal zipper motif (coiled-coil motif of DmKHC, *D. melanogaster*) for constitutive dimerisation and with two point mutations at amino acids 409 and 601. Cysteins were introduced at these positions for site-specific labelling. This construct is then named *Hsp90z_E409C_S601C*. The positions for labeling were chosen to ensure a stable FRET signal, unaffected by the large conformational rearrangements between the two monomers [26]. Theoretical FRET efficiency values are obtained from accessible volume simulations and distance calculations, done with the FRET positioning and Screening (FPS) software [27]. The calculations are described in Supplementary Note 1 and the results are summarized in Supplementary Tables 1 and 2. Hsp90 was expressed and purified as described in Supplementary Note 2 and then labeled via cysteine-maleimide chemistry with Lumidyne dyes (LD555 and LD655) as described in Supplementary Note 3. It is to be noted that specific dual labeling of Hsp90 is achieved via monomer exchange with a 20-fold excess of wild type Hsp90.

### Transfection of Biomolecules

Microinjection or a transfection with the toxin Streptolysin-O (SLO), were used to introduce labeled molecules in living HeLa cells. These methods were applied as follows: For **microinjection**, cells were grown to confluency at 37°C in cell culture imaging dishes (µ-Dish 35 mm, high Grid-500 Glass Bottom, Ibidi). Then esDNA was microinjected into living cells and then fixed with Para-formaldehyde solution (4%). The injection was monitored with a fluorescence microscope (Axio Observer 3, Carl Zeiss Microscopy GmbH., Oberkochen) equipped with a DIC module, a Colibri 7 LED light source, beam splitters, and a 20x Zeiss air objective. The injection system Femtojet 4i (Eppendorf) was used in combination with the micromanipulator InjectMan 4 (Eppendorf). Prior to injection, the cells were washed twice with pre-heated DPBS and then Opti-MEM was added. The samples were diluted in H1 buffer (40 mM HEPES, 150 mM KCl, 10 mM MgCl_2_, pH = 7.4). The injection needles Femtotip II (Eppendorf) were filled with the protein sample using microloader pipette tips. Injections were performed with the following settings: injection pressure=300 hPa, time=0.5 sec and compensation pressure = 30 hPa. Post-injection, Opti-MEM is removed and cells washed twice with DBPS. For fixation 500 µl of 4 % Paraformaldehyde (PFA) are added. Then the dish is incubated for 20 min at room temperature. For injection volume calibration and the number of injected molecules see Supplementary Note 4. **Streptolysin-O transfection** was done according to ref. [17]. HeLa cells were seeded in 8-well dishes one day prior to the experiment. The cell number was adjusted to achieve 90-100% confluency on the day of the experiment. The samples were diluted with degassed Tyrodes buffer (TB; 140 mM NaCl, 5 mM KCl, 1 mM MgCl2, 10 mM Hepes, 10 mM glucose, pH 7.4) to a final concentration of 400 nM and SLO was activated with TCEP. For this, SLO (25000 U/ml) and TCEP (final concentration 10 mM) were mixed and incubated for 20 min at 37 °C. For the transfection, the cells were first washed with DPBS and then incubated with a diluted SLO solution (125 U/ml in DPBS 1 mM MgCl_2_) at 37°C for 10 min. During incubation, SLO is allowed to insert into the plasma membrane and assemble to large pores. After incubation, the cells were gently washed with DPBS 1 mM MgCl_2_ and then incubated with the prepared sample for 5 min on ice. Following, the cells were washed with TB. For recovery, the cells were then incubated for 20 min with recovery medium (DMEM, 10% FBS, GlutaMAX [1x], sodium pyruvate [1x], 2 mM ATP) at 37 °C. To minimize background fluorescence, the cells were then washed with DPBS and finally imaged in live-cell imaging medium. All buffers and media were prewarmed at 37 °C.

### HILO Measurements

We used a custom HILO microscope (Supplementary Note 5 and Supplementary Fig. 2) where the observation depth could be controlled independently from the excitation beam, as other groups have done before [28]. Four excitation lasers were included for various measurements. The respective fluorescence can be detected in alternating laser excitation (ALEX) mode and result in three fluorescence signals as shown in Fig. 1c. The correction factors were determined as detailed in Supplementary Note 5. Analysis of the fluorescence signals involved co-tracking of molecules in all three channels (*Dem*|*Dex, Aem*|*Dex, Aem*|*Aex*). Here, *Dem Dex* is donor emission upon donor excitation, *Aem*|*Dex* is acceptor emission upon donor excitation and *Aem Aex* is acceptor emission upon acceptor excitation. This methodology enables accurate tracking even when conformational changes and therefore changes in FRET efficiency occur[29].

**Figure 1:**
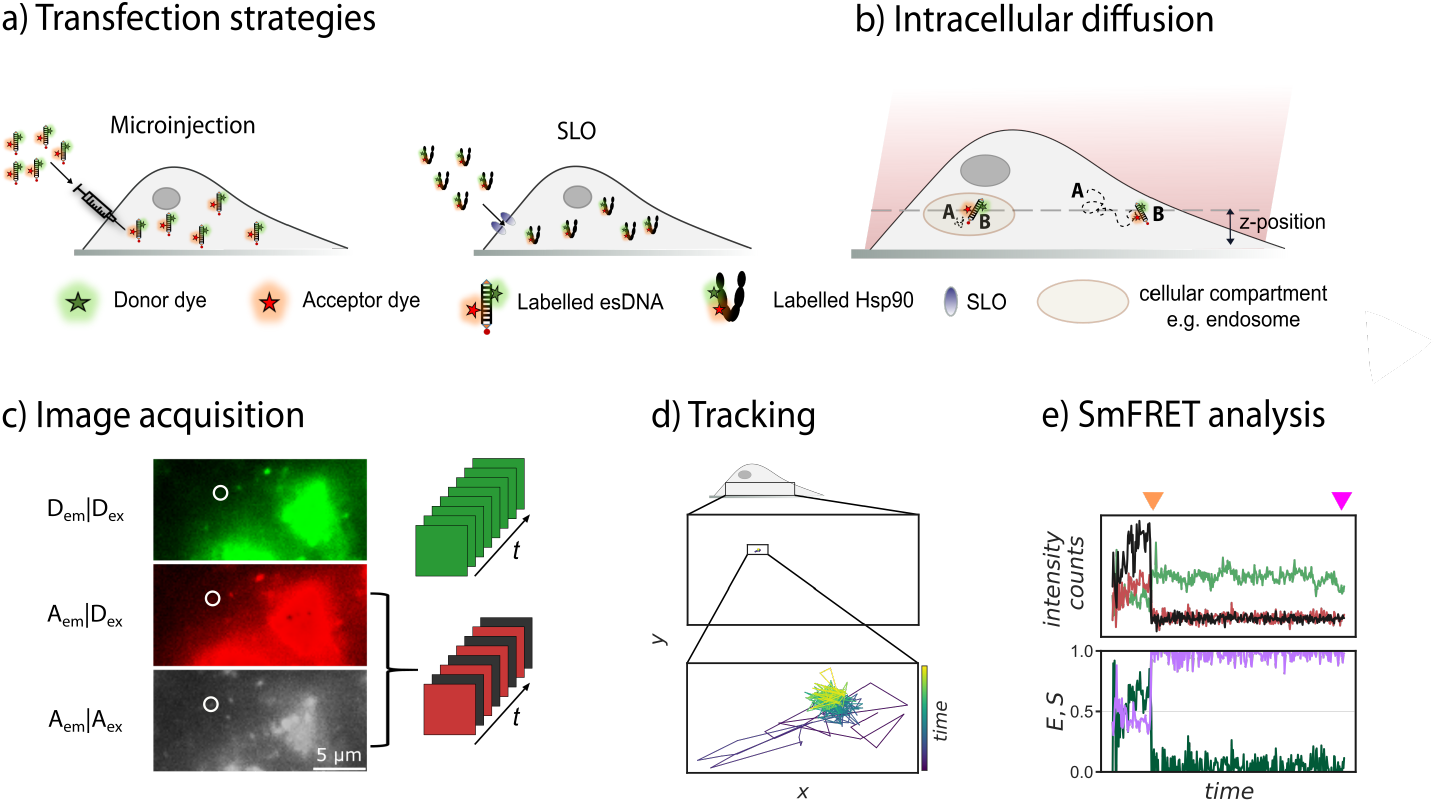
a) Schematic representation of microinjection (left) and Streptolysin-O transfection (right) of living HeLa cells. Tested biomolecules are end-sealed DNA and the protein Hsp90_E409C_S601C. b) Labeled biomolecules are tracked and three fluorescence signals are extracted from alternating laser excitation (ALEX) with a custom built HILO microscope. c) One frame of a SLO-transfected Hsp90 sample, demonstrating the ALEX excitation-emission strategy to obtain the three spatially resolved fluorescence signals *I*_Dem|Dex_; *I*_Aem|Dex_; *I*_Aem|Aex_. d) Example paths from a SLO-transfected Hsp90 protein tracked for 62 seconds. e) Example trace from the same protein showing *I*_Dem|Dex_ in green, *I*_Aem|Dex_ in red and *I*_Aem|Aex_ in black. The two arrows indicate acceptor and donor bleaching, respectively. All biomolecules were labeled with Lumidyne dyes LD555 (donor dye) and LD655 (acceptor dye), respectively. See main text for more details.

### Confocal Measurements

Single-molecule fluorescence experiments for Hsp90 *in vitro* were conducted at 295 K on a custom-built confocal setup as described before [30]. Green (532 nm, LDH-D-FA-530L, PicoQuant) and red laser light (640 nm, LDH-D-C-640, PicoQuant) were used to excite donor and acceptor dyes in pulsed interleaved excitation (PIE) mode [31] with an repetition rate of 20 MHz. The laser power is 317 µW for green and 114 µW for red, respectively, and was measured before the objective. Before focusing on the sample by a 60× water immersion objective (CFI Plan Apo VC 60XC/1.2 WI, Nikon), both beams were polarized and superimposed by a dichroic mirror (F43-537 laser beam splitter z 532 RDC, AHF). Another dichroic mirror (F53-534 Dual Line beam splitter z 532/633, AHF) separates excitation and emission light. In the emission path a third dichroic mirror (F33-647 laser beam splitter 640 DCXR, AHF) separated donor fluorescence from acceptor fluorescence. Pinholes (50*µm* diameter) filtered off-focus light. Before detection, polarizing beam splitters separated parallel and perpendicular polarized light. Donor and acceptor emission was detected after passing IR-filters (C-HSP750-25, LaserComponents) by single-photon detectors (two SPCM-AQRH-14-TR, Excelitas and two PDM series APDs, Micro Photon Devices).

The measurements were performed in SiPEG surface-passivated chambers. Therefore, high-precision coverslips (Carl Roth, 24 nm × 60 mm, (170 ± 5)*µm*) were cleaned by sonication in 2% Hellmanex (Hellma Analytics), ultra pure water and isopropanol: water mixture (1:3) to dissolve impurities. Remaining droplets were removed with nitrogen. The slide surface was activated by low pressure plasma (Tetra 30, Diener Electronics) performing subsequently an O_2_ and air plasma process. The plasma process was directly followed by treating the activated coverslips with methoxy-silane-polyethylenglycol (mSiPEG) solution (5 kDa, Rapp Polymere GmbH) with a concentration of 30 mg/mL. Unbound mSiPEG was washed away with pure water. The cavity for the single-molecule experiments were formed by a two-component silicon polymer (Microset 101, Microset Products Ltd). Before measurement, the chamber was preincubated with BSA (0.5 mg/mL) for about 20 min to prevent proteins from sticking.

The monomer-exchanged sample was diluted in degased Tyrodes buffer to a final concentration of 200 pM and incubated for about 6 h with 2 mM AMP-PNP. Microtimes and macrotimes of detected photons in each detection channel were recorded for 3600 s in T3 mode using time-correlated single-photon counting (HydraHarp400, PicoQuant), with temporal resolutions of 16 ps and 50 ns, respectively.

## Data Analysis

### Single-molecule experiments on HILO microscope

As the transfected molecules can freely diffuse within the cytosol, tracking of the molecules must precede the actual smFRET data analysis. Localisation and tracking of single particles was performed with a python-based software with Jupyter notebook interfaces, previously described in [32][29]. Upon tracking, the fluorescence intensity over time is extracted in the three channels *Dem*|*Dex, Aem*|*Dex* and *Aem*|*Aex* for each particle. With a generic user interface each recognized particle can be inspected. Based on fluorescence time traces, ES plots and the original movie a first manual selection of particles can be done. Only particles that show single-molecule features, such as single-step dye bleaching, extinction of the FRET signal upon bleaching of the first dye and anti-correlated signal upon acceptor bleaching, are selected. Traces with multiple dyes are not further analyzed. The selected traces are then saved in each one txt file. The txt files are imported into Igor Pro for further smFRET analysis as e.g. described in [33]. Importantly, in Igor Pro the traces are manually segmented in three parts: (1) prior to bleaching, when both dyes are present and FRET can happen, (2) after the first dye bleached and only one dye shows a signal, and (3) after the second dye bleached and only background signal remains. For further FRET analysis only the first part is used while parts two and three can be used for data correction.

The FRET efficiency *E* is calculated from the fluorescence intensities of the dyes and the correction factors with equation 1 for each point in time [25].

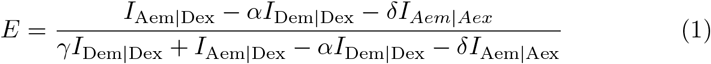

The stoichiometry *S* is calculated accordingly with equation 2.

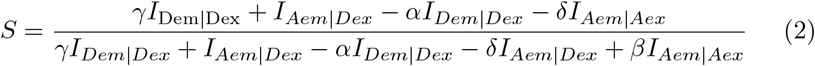

All FRET efficiency data is cumulated in histograms for a statistical analysis of the efficiency distributions. The correction factors *α* (crosstalk), *β* (differences in absorption cross-section and excitation powers of donor and acceptor fluorophores, respectively), *γ* (differences in quantum yield and detection efficiency) and *δ* (direct acceptor excitation) are given in Supplementary Table 6.

### Single-molecule experiments on confocal microscope

The analysis of the confocal data was done using the Wolfram Mathematica package Fretica(https://schuler.bioc.uzh.ch/programs) [34]. The ‘route correction matrix’ (RCM) corrects the number of photons for each detection channel. RCM was set by applying the correction factors according to the benchmark study [25]. The G-factor was assumed to be 1. Using the Δ*T* burst search algorithm with dual channel burst search (Fretica) with parameters Δ*T* = 150 µs, *N*_max_ = 100000, *N*_min_ was determined from the mean photon rate per second of the calculated background signal. The FRET efficiency *E* and the stoichiometry *S* were calculated for each identified single-molecule event according to equations 1 and 2, correction factors are given in Supplementary Table 6. Only single-molecule FRET events were taken into account in the Transfer Efficiency *E* histogram in figure 4a. Five Gaussian shapes with a constrained width of 0.09 were assumed as the fitting model. Please note that the population highlighted in gray is most likely caused by the strong donor-only populations mainly caused by single- or overlabeled Hsp90 proteins.

## Results and Discussion

### Benchmarking *in cellula* smFRET with esDNA

As cells inherently show high, inhomogeneous autofluorescence, we first wanted to understand whether the signal-to-noise ratio (SNR) of live-cell fluorescence time traces is sufficiently high to detect single-molecule FRET and to which extent the determined FRET efficiencies match with *in vitro* data. Therefore, we tested a stable and easy-to-transfect DNA sample with known dye distance, previously used for smFRET benchmark studies [25], labeled with lumidyne dyes LD555 and LD655. We compared the quality of the fluorescence time traces as well as the obtained peak FRET efficiencies in three different environments with increasing complexity: *in vitro*, in fixed-cells and in living cells (fig. 2), top panel. *In vitro* a high SNR with clear acceptor and donor bleaching can be obtained (fig. 2 a). Because the molecules are immobilized, they can be traced for extended periods of more than a minute. The average FRET efficiency from about 180 traces is *E* = 0.80, which replicates the values determined before [25]. In both, fixed and living cells, the SNR substantially decreases, leading to a reduced number of clearly identifiable single molecules. This is because fluctuations in background noise during the molecules’ diffusion are superimposed on the fluorophore signal. Nonetheless, a high SNR and clear bleaching steps can be obtained for a subset of traces (10-20 traces per dataset). Examples are shown in fig. 2b,c. With average values of 0.63 and 0.61 for fixed and live-cells respectively, the FRET efficiency decreases by approximately 25% compared to the efficiency determined *in vitro*. Considering the complex nature of the cellular environment, this behavior is not unexpected and may arise from multiple factors like dynamic quenching of the fluorophores caused by collisions with small molecules (e.g. oxygen) or by static quenching through interactions of the fluorophores with proteins binding to the DNA. Such effects are much more prominent in the macromolecular crowded cytosol and reduce the fluorescence intensity and therefore likely the FRET efficiency. The unambiguous smFRET signal (single donor and acceptor bleaching steps and anti-correlated behavior) proves that with our FRET-TTB strategy, we are able to i) track single molecules, ii) extract fluorescence intensities and iii) determine smFRET efficiencies and stoichiometries of labelled molecules in fixed and living cells.

**Figure 2:**
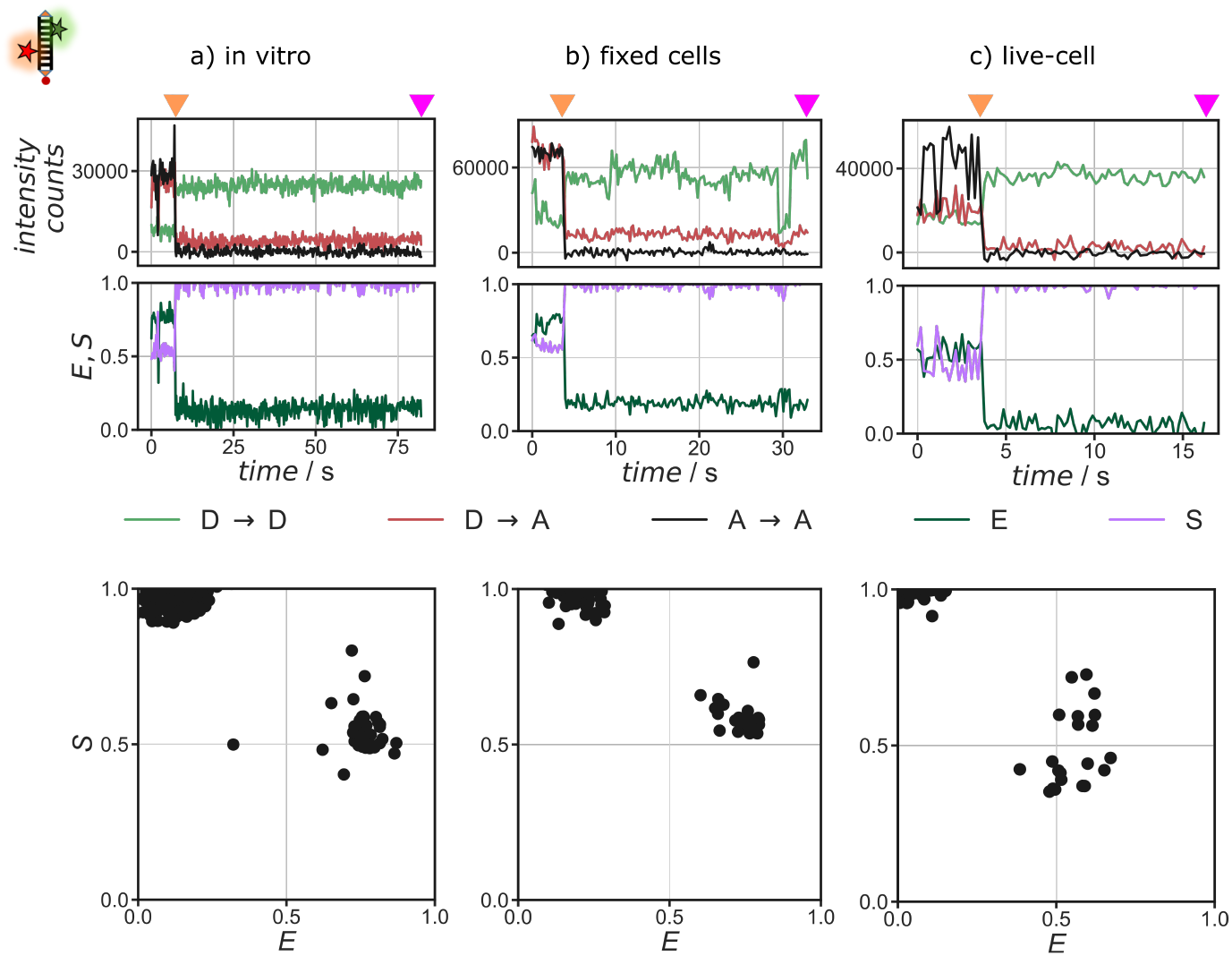
Comparison of *in vitro* and *in cellula* smFRET data for esDNA. a) in vitro; b) fixed-cell (microinjected) and c) live-cell (SLO transfection) data. Intensities *I*_Dem|Dex_ (green), *I*_Aem|Dex_ (red) and *I*_Aem|Aex_ (black) from tracked single molecules as well as their FRET efficiency (dark green) and stoichiometry (violet) against time. In addition, the raw FRET efficiency vs stoichiometry plots are shown at the bottom.

For further experiments with protein samples, we focused on the transfection with SLO. Microinjection is a low-throughput method and special instrumentation as well as extensive training are needed for successful transfections. This currently renders the method less efficient and less suited for a broad application.

### *In cellula* smFRET with the protein Hsp90

Next, we show that the transfection of HeLa cells also works with a dimer of Hsp90. Generally, complex multidimensional motions within proteins lead to a less stable fluorescence signal when compared to the rather stable, linear, short DNA oligos, used as a FRET-standard sample. While immobilization and background correction help maintain a high SNR *in vitro*, it is not possible for freely diffusing biomolecules in living cells. Consequently, many of the traces become difficult to interpret. By acquisition of large data sets, we are now able to obtain traces with a sufficient SNR for smFRET analysis. Clear single-step bleaching of the acceptor in fig. 3 indicates that single molecules are observed, while the anticorrelated increase in donor intensity is a strong indicator for FRET. Examples are shown in fig. 3b,c (left panel). Furthermore, the stoichiometry (*S*) versus FRET efficiency (*E*) plots (fig. 3b,c, middle panel) clearly separate a donor-only population at *S ≈* 1 and *E ≈* 0 and a FRET population at *S ≈* 0.5. Besides smFRET information, our analysis also provides access to tracking data, enabling us to monitor the intracellular movement of molecules. The two particles, shown in fig. 3 were tracked for about 62 s (792 frames) and 29 s (326 frames), respectively. From the corresponding trajectories (see fig. 3) diffusion coefficients of *D* = (5.6 ± 22) · 10^*−*6^*µ*m^2^*/*s and *D* = (1.1 ± 0.47) · 10^*−*4^*µ*m^2^*/*s were determined. Diffusion coefficients previously measured for various proteins in the cytosol of living prokaryotic and eukaryotic cells typically range from 10^*−*2^ − 10^2^*µ*m^2^*/*s [35]–[37] which is several orders of magnitude higher than the coefficients determined in our study. This indicates that instead of free diffusion largely non-diffusive, stagnant states have been observed. As Hsp90 does not have any anchor to restrict movement *in cellula*, it is likely that the observed molecules are trapped within cellular compartments such as endosomes or lysosomes. This work lays the groundwork for a more sophisticated analysis to, e.g., extract diffusion states [38]. By combining tracking and smFRET information, it will be possible to address many questions, such as whether the subcellular localization in different compartments can influence the conformational state.

**Figure 3:**
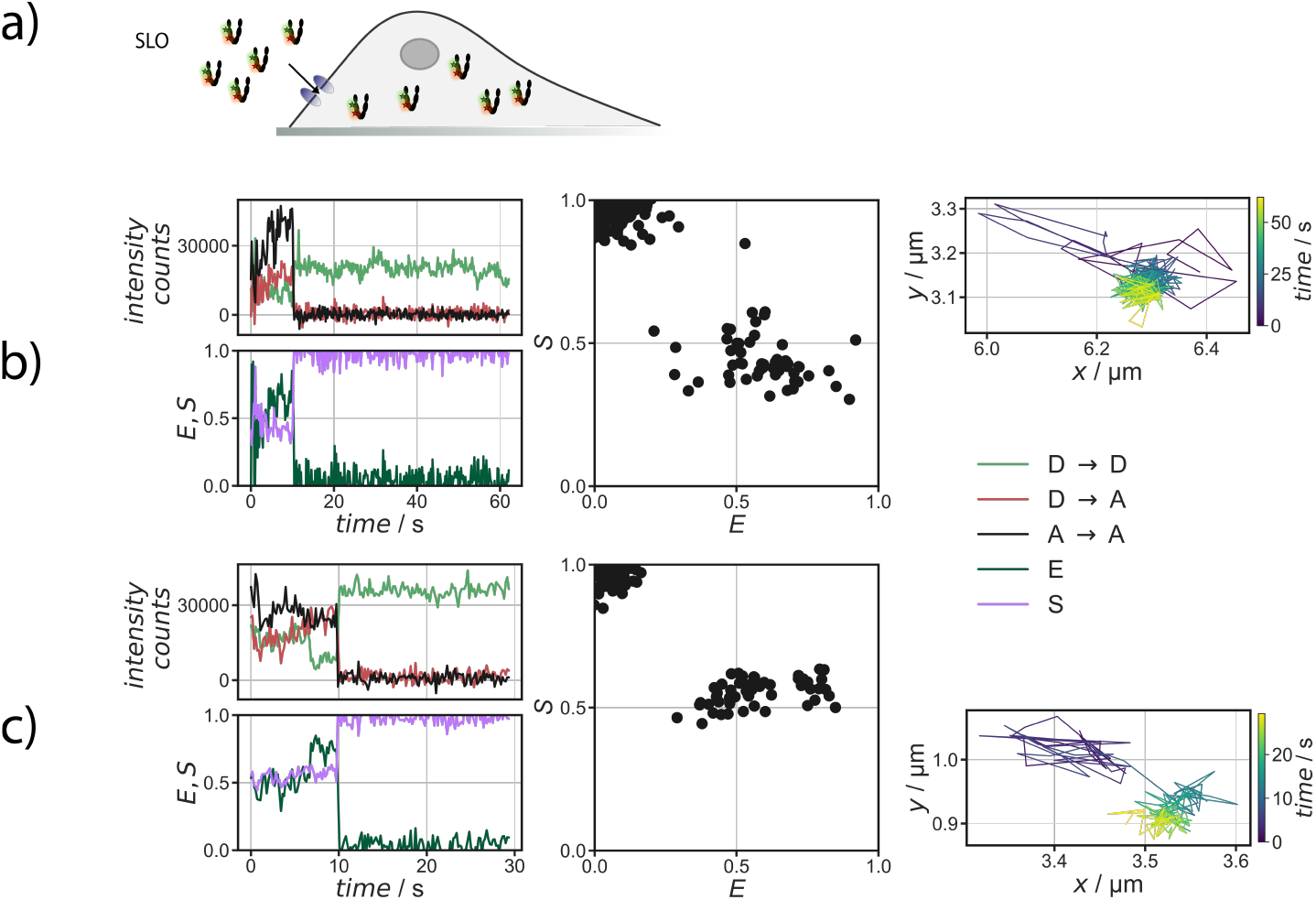
Examples of living cell single molecule fluorescence and FRET data. a) Schematics of the experiment with double labelled Hsp90 with LD555 and LD655 in cellula introduced via Streptolysin-O transfection. b,c) Two examples of fluorescence intensity traces, their corresponding stoichiometry vs FRET efficiency plots and tracks of these Hsp90 proteins (from left to right).

### Comparison of Hsp90’s conformational states *in vitro* and in living cells

Fig.4 shows a comparison of the FRET efficiencies obtained for Hsp90 *in vitro* and in living cells transfected with SLO. The overall agreement is remarkably good taking into account the different environment of these measurements. Fig.4a shows the results of measurements performed *in vitro*in the presence of 2mM AMP-PNP to populate the closed conformations of Hsp90, while Fig.4b presents the results of measurements taken within living cells. When we designed these labeling positions, we expected only two peaks at *E* = 0.354 and *E* = 0.435 for the open and closed Hsp90 conformations, respectively. This is because we exchanged the labeled Hsp90 with 20 times unlabeled Hsp90 before transfection, i.e., almost all Hsp90 dimers should only have two dyes in one monomer and no dyes in the other monomer (see Supplementary Note 1 for details). Indeed, both FRET efficiency distributions show large contributions at the expected FRET efficiency of around 0.4, but there are several additional peaks. These other FRET efficiencies can have several reasons. First, when the monomers are not fully exchanged and dyes are bound to both monomers of Hsp90, a multitude of FRET efficiencies can be expected (see Supplementary Tables 1 and 2 for all possible combinations). Second, the labeling position 601 is not resolved in the yeast Hsp90 crystal structure (2cg9) and therefore is likely quite flexible. Third, the many interactors, such as cochaperones, clients in the cell might affect the distance between amino acids 409 and 601, causing different FRET efficiencies. For a quantitative comparison, we chose to fit four Gaussians to the FRET efficiency distributions *in vivo*, and an additional one for excluding artefacts most likely from single- or double labeled protein samples *in vitro*. The widths of all Gaussians were fixed at 0.09. Table 1 shows a comparison of these four most pronounced contributions. Altogether, we see a remarkable agreement for the peak positions for Hsp90’s states with this FRET pair *in vitro* and *in vivo*. Some differences are expected, given the very crowded environment of the cell and possibly many specific interactions with other proteins in the cell. This result again underlines the applicability of our FRET-TTB approach to living cells and opens now the path to investigate proteins in their native environment.

**Figure 4:**
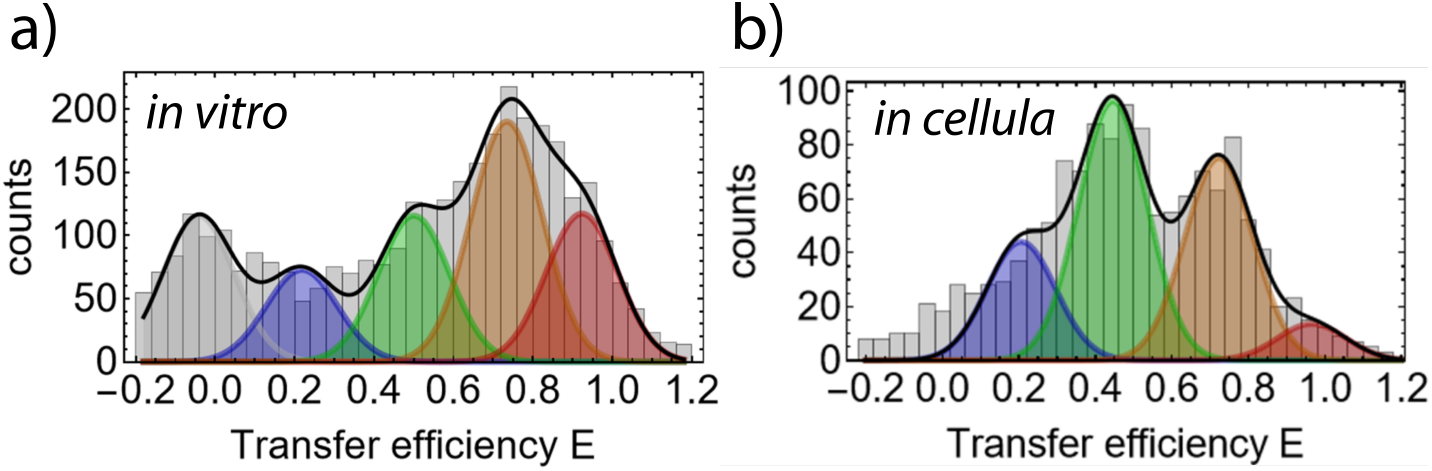
Comparison of *in vitro* (a) and *in cellula* (b) FRET efficiencies for Hsp90. *In vitro* smFRET-PIE data was recorded on a confocal microscope for 1 hour, meausured in presence of AMP-PNP. *In cellula* data from 51 traces on the HILO microscope. Four Gaussian were fit to the data to compare the two data sets. For the confocal data a additional very low FRET Gaussian was used (gray), which accounted for bursts most likely from single (donor only) labeled proteins.

**Table 1:** Comparison of the *in vitro* and *in cellula* FRET efficiency distributions for yHsp90_E409C_S601C. E - mean efficiency of the Gauss fits, occ. - relative abundance of FRET efficiencies within the Gauss fits.

|  | *in vitro* |  | *in cellula* |  |
| --- | --- | --- | --- | --- |
|  | *E* | occ. /% | *E* | occ. /% |
| low FRET | 0.21 | 15 | 0.21 | 19 |
| mid FRET 1 | 0.50 | 23 | 0.45 | 42 |
| mid FRET 2 | 0.73 | 38 | 0.72 | 33 |
| high FRET | 0.92 | 24 | 0.97 | 6 |

## Conclusion

The approach we developed provides a controlled and effective means of observing and understanding the multidimensional dynamics of biomolecules within living cells. This is essential in order to gain insight into the functions of proteins in various cellular compartments. We demonstrated the power of this method by first establishing and validating a robust strategy using end-sealed DNA (es-DNA) to achieve single-molecule Förster Resonance Energy Transfer (smFRET) in live cells. We then applied this technique to investigate the behavior of the Hsp90 protein in a living cellular environment, revealing results that were consistent with the *in vitro* data obtained from confocal smFRET experiments. With our home-built Highly Inclined and Laminated Optical sheet (HILO) setup, we were able to observe and track single DNA and protein molecules several microns into the cell cytoplasm and other compartments. Alternating laser excitation (ALEX) helped to obtain stoichiometry information and to clearly distinguish states with different FRET efficiencies. The alternating laser excitation strategy was crucial to obtain signal information in three channels and therewith to access single-molecule stoichiometry information apart from known photo-physical properties of dyes *in cellula*, which otherwise might be misinterpreted as dynamics. In conclusion, we have shown a method based on single-molecule FRET tracking and transfection of biomolecules (FRET-TTB) to investigate single biopolymers in living cells. FRET-TTB has enabled us to step into long-term fluorescent intensity analysis with smFRET and stoichiometric information, which was not possible before in the cytosol of cells due to technical limitations, short observation times and short-lived fluorescent proteins. Furthermore, the developed method is not restricted to a specific area of the cell and is therefore a valuable tool for investigating protein plasticity and localization-dependent function of proteins. The precise control of the transfected proteins enables us to quantify the response of biomolecules to different cellular environments or compartments in an unprecedented way. Our study can be further used to bridge the gap between *in vitro* and *in cellula* measurements.

## Acknowledgements

This work was supported by the Deutsche Forschungsgemeinschaft (DFG) under Germany’s Excellence Strategy (CIBSS EXC-2189 Project ID 390939984) and the SFB1381 (Project ID 403222702) and by the European Research Council through ERC grant agreement no. 681891. We thank Dr. Fernando Aprile-Garcia and Dr. Ritwick Sawarkar for help with setting up the cell culture and the viability assays, we thank Dr. Bianca Hermann and Jolanta Vorreiter and Michael Witt for their help in protein production of Hsp90 variants, we thank Marianne Birkle for help with the cell cultures, we thank Prof. Dr. Ben Schuler for aiding us in running our initial tests of Hsp90 on their confocal injection setup, we thank Dr. Bismark Appiah for providing his expertise in handling the Femtojet II injector and micromanipulator for cell experiments, we thank Dr. Yasser Gidi and Dr. Liwei Zheng for helpful discussions. ChatGPT, developed by OpenAI, assisted by providing some suggestions for language improvements in some paragraphs.

## Author Contributions

A.A. and T.H. designed research; A.A. performed the microinjection experiments and data analysis; A.E. performed the SLO experiments and data analysis. V.F. performed the confocal microscopy experiments and data analysis. P.W., A.A performed the initial tests with SLO; L.S. and G.J.S. provided and optimized the three-channel tracking software with the help of A.A. and A.E.; A.A, T.H and C.L. designed and co-built the HILO setup; A.A., A.E. and T.H. wrote the manuscript with the help of all authors. All authors have approved the final version of the manuscript.

