## Supplementary Notes 1-6 for "Single-Molecule FRET and Tracking of Transfected Biomolecules in Living Cells"

### Supplementary Information for: Single-molecule FRET and Tracking of Transfected Biomolecules in Living Cells

1. **Suppl. Note 1: FRET efficiency calculations** 2
2. **Suppl. Note 2: Protein production** 5
3. **Suppl. Note 3: Labelling of Biomolecules** 6
4. **Suppl. Note 4: Microinjection** 7
5. **Suppl. Note 5: Tracking Microscopy (HILO)** 9
6. **Suppl. Note 6: Data Analysis** 10

*Equal contribution

#### Supplementary Note1: AV and distance calculation

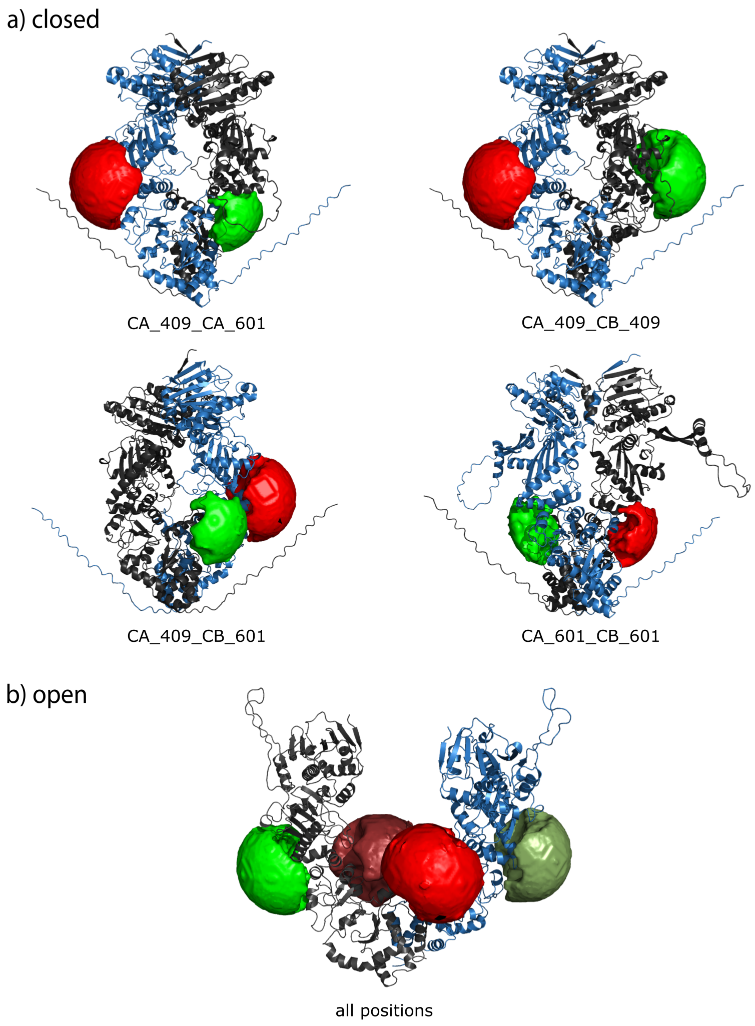


Figure 1: Some possible combinations of labeling positions for the cysteins at positions 409 and 601. Upper four figures based on an Alphafold2 structure for closed yeast Hsp90. Bottom based on the open structure of yeast Hsp90 [1]. Accessible volumes for donor and acceptor are based on FPS analysis [2]. They are depicted in green and red, respectively. CA: Chain A; CB: Chain B.

Accessible dye volume (AV) simulations and distance calculations were performed with the Lumidyne dye pair LD555 and LD655 (Lumidyne Tech). As the original crystal structure of Hsp90 (2cg9) does not fully resolve the C-terminal end, we used the Alphafold2 structure for closed yeast Hsp90. The open structure of Hsp90 (m-om4) was published before in [1]. Assuming that exactly two dyes are bound to Hsp90, the distances for all possible combinations of dye positions were calculated using the software FPS [2]. The AVs for various combinations are depicted in Suppl.Fig. 1 and the calculated distances and FRET efficiencies are summarized in Suppl. Table 1.

Table 1: Distances for different combinations of labeling positions for the closed Hsp90_409_601 dimer. Calculations were done with dye parameters for LD555 and LD655. Specifications of the two dyes (#1, #2): the dye-position (CA-chain A; CB-chain B) and the kind of dye (A-Acceptor; D-Donor); Rmp-distance between the mean dye positions; $\langle RDA\rangle E$-FRET averaged distance; E-FRET Efficiency.

| #1 | #2 | $Rmp$ | $\langle RDA\rangle$ | $sigmaDA$ | **E** | $\langle RDA\rangle E$ |
|---|---|---|---|---|---|---|
| CA_409_A | CA_601_D | 63,7 | 65,3 | 7 | 0,435 | 64,8 |
| CA_409_A | CB_409_D | 82,8 | 84,7 | 7,1 | 0,147 | 83,2 |
| CA_409_A | CB_601_D | 33,5 | 36,3 | 8,2 | 0,937 | 39,5 |
| CA_409_D | CA_601_A | 63,7 | 65,3 | 7 | 0,435 | 64,8 |
| CA_409_D | CB_409_A | 82,8 | 84,7 | 7,1 | 0,146 | 83,2 |
| CA_409_D | CB_601_A | 33,5 | 36,3 | 8,2 | 0,937 | 39,5 |
| CA_601_A | CA_409_D | 63,7 | 65,3 | 7 | 0,435 | 64,8 |
| CA_601_A | CB_409_D | 33,5 | 36,2 | 8,1 | 0,938 | 39,4 |
| CA_601_A | CB_601_D | 48,8 | 49,8 | 5,9 | 0,771 | 50,6 |
| CA_601_D | CA_409_A | 63,7 | 65,3 | 7 | 0,435 | 64,8 |
| CA_601_D | CB_409_A | 33,5 | 36,2 | 8,1 | 0,938 | 39,4 |
| CA_601_D | CB_601_A | 48,8 | 49,8 | 5,9 | 0,771 | 50,6 |
| CB_409_A | CB_601_D | 63,8 | 65,4 | 7 | 0,434 | 64,8 |
| CB_409_A | CA_409_D | 82,8 | 84,8 | 7,1 | 0,146 | 83,2 |
| CB_409_A | CA_601_D | 33,5 | 36,3 | 8,1 | 0,938 | 39,5 |
| CB_409_D | CB_601_A | 63,8 | 65,4 | 7 | 0,434 | 64,8 |
| CB_409_D | CA_409_A | 82,8 | 84,8 | 7,1 | 0,146 | 83,2 |
| CB_409_D | CA_601_A | 33,5 | 36,3 | 8,1 | 0,938 | 39,5 |
| CB_601_A | CB_409_D | 63,8 | 65,4 | 7 | 0,434 | 64,8 |
| CB_601_A | CA_409_D | 33,5 | 36,3 | 8,2 | 0,937 | 39,5 |
| CB_601_A | CA_601_D | 48,8 | 49,9 | 5,9 | 0,771 | 50,6 |
| CB_601_D | CB_409_A | 63,8 | 65,4 | 7 | 0,434 | 64,8 |
| CB_601_D | CA_409_A | 33,5 | 36,3 | 8,2 | 0,937 | 39,5 |
| CB_601_D | CA_601_A | 48,8 | 49,9 | 5,9 | 0,771 | 50,6 |

Table 2: Distances for different combinations of labeling positions for the open Hsp90_409_601 dimer. Calculations were done with dye parameters for LD555 and LD655. Specifications of the two dyes (#1, #2): the dye-position (CA-chain A; CB-chain B) and the kind of dye (A-Acceptor; D-Donor); Rmp-distance between the mean dye positions; $\langle RDA \rangle E$-FRET averaged distance; E-FRET Efficiency.

| #1 | #2 | $Rmp$ | $\langle RDA \rangle$ | $sigmaDA$ | **E** | $\langle RDA \rangle E$ |
|---|---|---|---|---|---|---|
| CA_409_A | CA_601_D | 68,1 | 70,2 | 9,9 | 0,354 | 68,6 |
| CA_409_A | CB_409_D | 111,9 | 113,4 | 7,8 | 0,029 | 111,5 |
| CA_409_A | CB_601_D | 47,5 | 50,6 | 8,7 | 0,742 | 52 |
| CA_409_D | CA_601_A | 68,1 | 70,2 | 9,9 | 0,354 | 68,5 |
| CA_409_D | CB_409_A | 111,9 | 113,3 | 7,8 | 0,029 | 111,5 |
| CA_409_D | CB_601_A | 47,5 | 50,6 | 8,7 | 0,742 | 52 |
| CA_601_A | CA_409_D | 68,1 | 70,2 | 9,9 | 0,354 | 68,5 |
| CA_601_A | CB_409_D | 47,1 | 50,3 | 8,8 | 0,747 | 51,8 |
| CA_601_A | CB_601_D | 33,8 | 37,4 | 12,2 | 0,9 | 43 |
| CA_601_D | CA_409_A | 68,1 | 70,2 | 9,9 | 0,354 | 68,5 |
| CA_601_D | CB_409_A | 47,1 | 50,3 | 8,8 | 0,747 | 51,8 |
| CA_601_D | CB_601_A | 33,8 | 37,4 | 12,2 | 0,9 | 43 |
| CB_409_A | CB_601_D | 67,8 | 69,8 | 9,8 | 0,359 | 68,3 |
| CB_409_A | CA_409_D | 111,9 | 113,3 | 7,8 | 0,029 | 111,5 |
| CB_409_A | CA_601_D | 47,1 | 50,3 | 8,8 | 0,748 | 51,7 |
| CB_409_D | CB_601_A | 67,8 | 69,9 | 9,8 | 0,359 | 68,3 |
| CB_409_D | CA_409_A | 111,9 | 113,3 | 7,8 | 0,029 | 111,5 |
| CB_409_D | CA_601_A | 47,1 | 50,3 | 8,8 | 0,748 | 51,7 |
| CB_601_A | CB_409_D | 67,8 | 69,9 | 9,8 | 0,36 | 68,2 |
| CB_601_A | CA_409_D | 47,5 | 50,6 | 8,7 | 0,742 | 52 |
| CB_601_A | CA_601_D | 33,8 | 37,4 | 12,2 | 0,9 | 43 |
| CB_601_D | CB_409_A | 67,8 | 69,9 | 9,8 | 0,359 | 68,3 |
| CB_601_D | CA_409_A | 47,5 | 50,6 | 8,7 | 0,742 | 52 |
| CB_601_D | CA_601_A | 33,8 | 37,4 | 12,2 | 0,9 | 43 |

#### Supplementary Note2: Protein production

**Hsp90z_Avi_409C_601C:** Cysteine residues were introduced into the sequence of yeast Hsp90 via site-directed mutagenesis at positions 409 and 601 (Quick Change Lighting kit, Agilent). The cysteine modified Hsp90 was expressed with a pET28 vector. Additionally this protein has an Avi-tag, which leads to a biotinylation in the bacterial cell with the help of biotin ligase enzymes co-expressed in a pBirAcm (Avidity Nanomedicine, La Jolla, CA) plasmid. The cells were grown in LB media supplemented with 0.5% glucose and Kanamycin (100µg/ml) at 37 °C. At a OD of 0.7, the expression was induced by adding isopropyl $\beta$-D-1-thiogalactopyranoside (IPTG; Sigma; final concentration 1 M). Furthermore, biotin (prepared in 10 mM Bicine Buffer pH 8.3 (adjusted with 1 M KOH, filter sterilized)) was added to the medium with a final concentration of 50 $\mu$M. The cells were then harvested 4 hours after induction and pelleted by centrifugation at 6000 rpm. Cell pellets are stored at $-20°$C until protein purification. For cell lysis pellet was resuspended in NiNTA A buffer with 20 mM imidazole. Protease inhibitor 50x is added prior to cell disruption in a cell disruptor (I & L Biosystems GmbH). $MgCl_2$ (final concentration 2 mM) and 6 µl of Benzonase (EMD Millipore Corp, USA) are added to the cell lysate. It is then centrifuged for 1 h at 50,000g. The supernatant is then passed through a 0.45 filter (Merck Millipore millex), the filtrate is then collected and fed to the HisTrap. The filtrate collected is then fed to the sample pump for HisTrap1. The NiNTA HisTrap column is pre-washed and equilibrated with HisA1 buffer. The sample is loaded onto the column and then allowed to run through the HisTrap1. The bound fraction is collected. The outlet collected from HisTrap1 is fed again into the HisTrap2. The bound fraction is again collected and diluted so the total salt concentration is less than 30mM. This sample is then fed into HiTrapQ. The eluted fraction is then collected and then concentrated to less than 3 g/l and injected into the Size exclusion column 26/600. The final purified samples are collected, concentrated and frozen in liquid nitrogen for later use.

#### Supplementary Note 3: Labeling of biomolecules

The protein is labelled based on cysteine maleimide chemistry. The protein is reduced with 1 mM Tris (2-carboxyethyl) phosphine (TCEP) pH 7.0 for 30 minutes at room temperature. Then the protein is equilibrated with ice cold phosphate saline buffer pH 6.7 by washing in a centrifugal 50kDa cut-off spin filter (3 times) which removes excess TCEP. Then 1-2 fold (mol) of dye (LD555 and LD655) is added. First the acceptor dye (LD655) is added and incubated for 2 minutes, then the donor dye (LD555) is added. The mix is incubated at room temperature for 2 h. The labelled protein is then purified with PD MiniTrap G25 columns (Cytiva). The column is washed thrice with Tyrode's buffer. Then the protein is added and the column centrifuged for 2 min, at 1000 g. The flow through was collected and used for the experiments.

To create heterodimers with one doubly labeled monomer and one unlabeled monomer, a monomer exchange was performed. The labeled protein is mixed with an unlabeled variant with a C-terminal zipper (Hsp90z_wt). A ratio of 1:20 (labeled:unlabeled) is used and final concentrations of 2 $\mu$M and 80 $\mu$M in Tyrode's buffer are used. The proteins are incubated at 43°C for 36 min with shaking at 300rpm in a thermocycler (Eppendorf). The heating destabilizes the C-terminal zipper and enables exchange of the monomers. Possible aggregates are removed subsequently by centrifugation at 14,000 g and 4 °C for 30 min.

#### Supplementary Note 4: Microinjection

Cell viability was tested by using pHrodo Red and Green AM intracellular pH indicator dyes (ThermoFisher Scientific). The cells used for injection were first calibrated for the dye response with different pH buffer conditions using the Intracellular pH calibration Buffer kit (ThermoFisher Scientific). Imaging of the cells was done on an Olympus microscope fitted with a HE90 dichroic cube. Images of the cell area of the different dishes were taken before dye addition. After addition of dye, washing and incubation with pH calibration buffers for the respective cell dishes an image of the same area was recorded. For cell viability test new cell dishes were taken and areas for injection were imaged in OptiMEM before injection. The following injection of cells were made with H1 buffer, then the cells were washed with preheated PBS (twice) and imaged in optiMEM. Images of the cell area of the different dishes was taken before dye addition. After addition of dye, washing and incubation with OptiMEM the respective areas of dishes were imaged. Uninjected area cells were also imaged as a control. For intensity measurements, mean of 10 cells were taken for each of the three time points (1st hour, 2nd hour, 3rd hour) for the 4 different pH measurements (pH 4.5, pH 5.5, pH 6.5, pH 7.5). For before and after measurements care was taken to measure the same cells and same region. All measurements were background corrected.

| **Parameter** | **Value** |
|---|---|
| Speed (Coarse-mode) (µm/sec) | 4000 |
| Step injection speed (µm/sec) | 50 |
| Step injection distance (µm/sec) | 20 |
| Position speed (µm/sec) | 1500 |
| Injection axial | ON |
| Piezoxp. Speed (µm/sec) | 300 |
| Piezoxp. Distance (µm) | 20 |
| Piezoxp. axial | OFF |
| Angle of injection | 45° |

Table 3: Injector Parameters

Fixed cell experiments: The purified DNA/protein sample of 1nM is prepared in a vial with degassed buffer which is double filtered. Injection needle is filled with 5 µl of 1nM sample and fitted to the injector nozzle. Before injection, the cell area to be injected is imaged in the DIC, Green (555nm), Red(630nm) emission channels of the fluorescence microscope to check for any leakage or fluorescence abnormalities. The grid number on the ibidi dish is noted. The cells are then washed twice with DPBS 1ml (pre-warmed at 37°C) and then 1 ml of Opti-MEM is added, then cells within the marked grid area are injected. After injections, cell area within the grid is imaged again with DIC, green and red channels to confirm no leakage. Then the Opti-MEM is removed, cells washed with DBPS twice followed by addition of 500 µl of 4 percent Paraformaldehyde

(PFA). The dish is incubated for 20 mins in dark at room temperature. Then the PFA is washed twice with in DPBS and 2ml of Opti-MEM was added. The cells were then ready to be imaged.

#### Supplementary Note 5: HILO microscopy

For all *in cellula* measurements a home-built HILO microscope as depicted in Suppl.Fig. 2 was used. A theoretical depth of view calculation was done to estimate the minimum frame requirement for a molecule to be visible with a 70 ms exposure time and a certain diffusion constant. For a 100x Nikon objective with a 1.49 NA, the theoretical depth of field is 190 nm. The calculated depth of field $d_{\text{tot}}$ as per the below parameters was found to be:

$$d_{\text{tot}} = \frac{\lambda n}{\text{NA}^2} + \frac{n \cdot e}{M \cdot \text{NA}} \tag{1}$$

Substituting given values:

$$d_{\text{tot}} = \frac{553\,\text{nm} \cdot 1.515}{1.49^2} + \frac{1.515 \cdot 14\,\mu\text{m}}{200 \cdot 1.49} = 449\,\text{nm} \tag{2}$$

Here, the refractive index $n = 1.515$; wavelength $\lambda = 553\,\text{nm}$; the variable $e = 14\,\mu\text{m}$ is the smallest distance that can be resolved by a detector placed in the image plane of the microscope objective, whose lateral magnification is $M = 200$. In the following we estimate the displacement of a single molecule (in z-direction) that can still be tracked, $x_{\text{tr}}$, assuming the molecule is first detected in the center of the focus. We start from the depth of field of the microscope as:

$$x_{\text{tr}} \approx \frac{1}{2} d_{\text{tot}} \tag{3}$$

Substituting (2) in (3), $x_{\text{tr}} \approx 224\,\text{nm}$ is the maximum possible mean squared displacement, where the molecule can be tracked, i.e., beyond this distance the particle might leave the focus and cannot be tracked. The mean squared displacement in one dimension, $x^2$, of a molecule is related to the diffusion coefficient $D$ for a time $t$ as:

$$t \approx \frac{x^2}{2D} \tag{4}$$

$$x \approx \sqrt{2Dt} \tag{5}$$

We can now calculate the mean squared displacement for a given diffusion constant and time. E.g. for $D = 0.05\mu\text{m}^2/\text{s}$ and t=10s we would obtain $x = 33$ nm. Therefore, most molecules would not leave the focus for such an experiment. **Correction factors** were estimated for esDNA and Hsp90_409C_601C. Both were labelled with LD655 and LD555 and both were biotinylated. The esDNA and Hsp90 were separately immobilized on PEG surfaces coated with Biotin-NHS PEG. Then the molecules were attached to the surface via biotin-Neutravidin binding and measured in H1 buffer pH 7.5 with Tris (5 mM Tris, 20 mM $MgCl_2$, 5 mM NaCl, pH 7.5) at 10pM for esDNA. The measurements were carried out with ALEX. The movies were analyzed using homewritten procedures based on IgorPro (wavemetrics), which is detailed in [3]. The obtained correction factors are given in Supplementary Table 6.

#### Supplementary Note 6: Data Analysis

The data analysis is done with a python based code called fret-analysis, previously described in [4] and available on GitHub. Analysis starts by running the Jupyter Notebook titled "01. Tracking". First, the microscope movies (TIFF file stacks) must be provided along with calibration files for the laser profile and channel alignment. Channel alignment is done prior to tracking. For this, threshold parameters for feature radius, mass and signal intensity need to be set so that fiducial markers (e.g. fluorescent beads) in the registration calibration file are recognized. The same procedure is repeated for the actual data files. Parameters should be chosen to locate single-molecule signals while minimizing the recognition of unspecific background fluorescence. The parameters used in this study are listed in table 4. For tracking the Crocker-Grier algorithm was chosen. Several tracking parameters can be adjusted (see table 5). The tracking results are saved into six separate files. Next, the notebook "02. Analysis AE" is executed. This notebook is based on the notebook "02. Analysis" provided in [4]. However, the built-in data correction is skipped (set $\alpha = 0, \delta = 0, \beta = 1, \gamma = 1$), because instead we employ our home written Igor Pro Software for correction. The updated data is again saved into six files. A manual trace selection is then carried out using the GUI tool "inspector_gui", which can be launched from the command line (python -m smfret_analysis.inspector_gui). This tool allows for visual inspection of each recognized and tracked particle. An overview of the following information is shown for each particle: its position in the three separate movies (donor, FRET and acceptor); intensity traces over time for each channel; and the SE plot. Based on this information, users can manually accept or reject traces for further analysis. Selections are saved by clicking the appropriate button in the interface. In the third step, the selected traces are exported as text files. This is done by running the adjusted notebook "03. Plots" up to the plot_summary function with the export="True" argument. These .txt files are subsequently used for further analysis in Igor Pro. To import the traces into Igor Pro, the function "Importer_ascii()" is used. To access the data the "SetNotes()" function must be called. The traces can then be viewed via the "show saved traces" panel. Each trace is then manually segmented into three regions: 1. Potential FRET – both dyes are present. 2. Single dye – one dye has bleached. 3. Background – both dyes have bleached. From this segmentation, SE 2D histograms and the corresponding 1D histograms can be generated. Correction can be done interactively. In this study, the correction factors were determined with in vitro measurements of esDNA, see Table 6.

Table 4: Localization and Tracking Parameters.

| Parameter | Donor | Acceptor |
|---|---|---|
| **Registration options** | | |
| Algorithm | Crocker-Grier | Crocker-Grier |
| Radius | 2 | 2 |
| Signal Threshold | 1000 | 500 |
| Mass Threshold | 1000 | 1000 |
| **Localization options** | | |
| Algorithm | Crocker-Grier | Crocker-Grier |
| Radius | 2 | 2 |
| Signal Threshold | 1000 | 500 |
| Mass Threshold | 20000 | 13000 |

Table 5: Linking of Particles Parameters.

| Parameter | Value |
|---|---|
| Feature radius (feat_radius) | 2 |
| Link memory (link_mem) | 20 |
| Minimum length (min_length) | 20 |
| Background frame (bg_frame) | 3 |
| Background estimator (bg_estimator) | mean |
| Lag time | infinity |

Table 6: Analysis Correction Factors.

| Correction factors | HILO | Confocal |
|---|---|---|
| Leakage $\alpha$ | 0.17 | 0.23 |
| Direct Excitation $\delta$ | 0.12 | 0.05 |
| Excitation Efficiency $\beta$ | 1 | 0.7 |
| Detection Efficiency $\gamma$ | 0.8 | 1 |

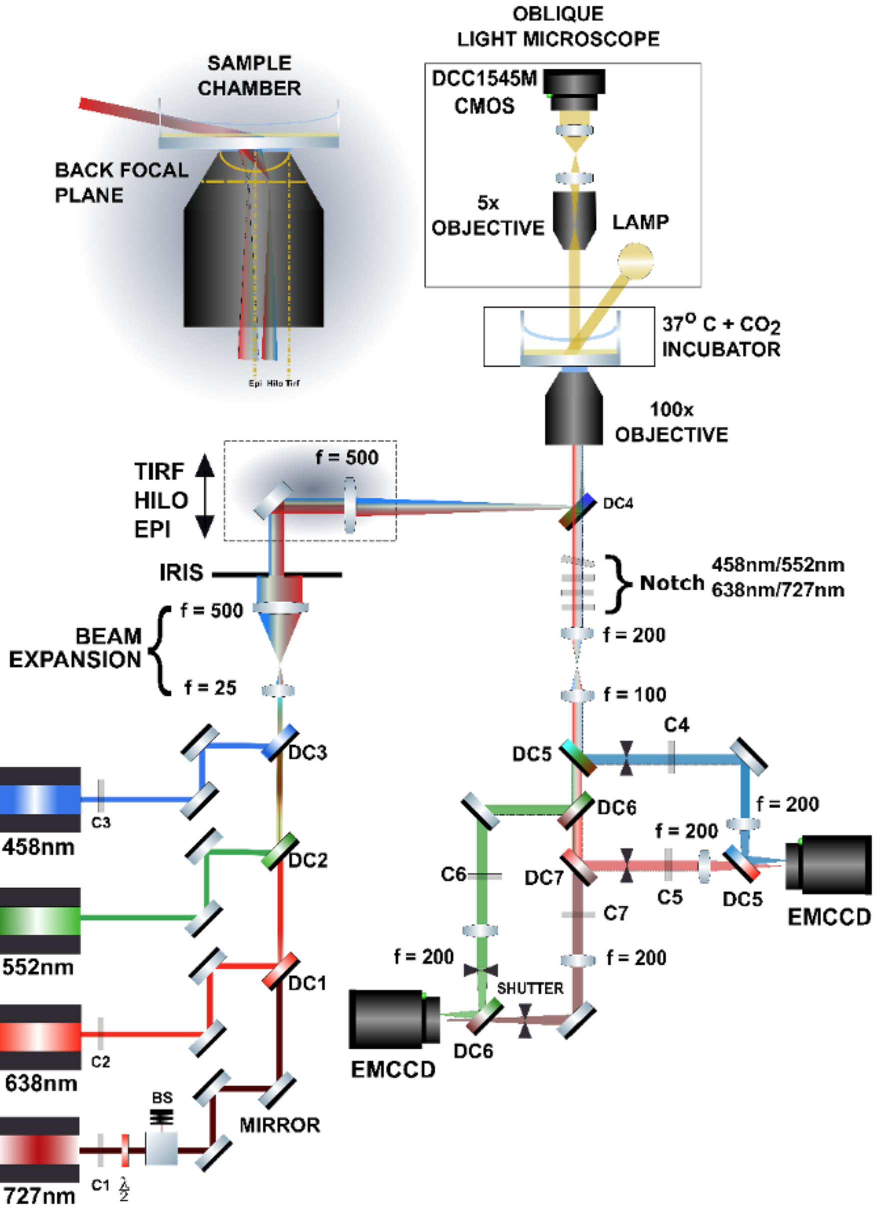
OBLIQUE
LIGHT MICROSCOPE
DCC1545M
CMOS
5x
OBJECTIVE
LAMP
SAMPLE
CHAMBER
BACK FOCAL
PLANE
37$^{O}$ C + $CO_2$
INCUBATOR
100x
OBJECTIVE
TIRF
HILO
EPI
f = 500
DC4
IRIS
Notch
458nm/552nm
638nm/727nm
f = 500
BEAM
EXPANSION
f = 25
f = 200
f = 100
DC3
DC5
C4
C3
458nm
DC6
f = 200
f = 200
DC2
C6
DC7
C5
DC5
EMCCD
552nm
C7
f = 200
f = 200
DC1
SHUTTER
C2
638nm
EMCCD
DC6
BS
MIRROR
727nm
C1
$\frac{\lambda}{2}$

Figure 2: Schematic representation of the 4-color HILO microscope equipped with an oblique light microscope for viewing cell injection area. All the optical and live-cell components are mounted on a vibration-damping optical table (RS2000, Newport). The following four lasers are used in our custom-built setup: 458nm (Obis 458LX, Coherent), 638nm (Obis 637LX, Coherent), 552nm (Obis 552LS, Coherent), 727nm (Obis 730LX, Coherent). The lasers are cleaned up using clearance filters and then overlaid using specific dichroic mirrors DC1, DC2 and DC3. An iris is positioned to select homogeneous illumination for all overlaid lasers. Subsequently the beam is expanded 20x by a Keplerian telescope and further a 2-inch iris cuts out the central illumination region. Further this beam is focused on to the back focal plane of the objective via a multiband Quad Line beam splitter DC4 using a 2-inch cylindrical lens mounted on a translation stage. This translation stage helps to shift the light path away from the optical axis (epi) to HILO and TIRF regions. The objective used is (CFI Apo TIRF 100x, Nikon) with NA=1.49. A set of notch filters are setup after the Quad Line beam splitter to block the excitation beam. The emission fluorescence is then directed to the detection box where the beam passes through a Keplerian lens with an adjustable optical slit (Owis GmbH). This helps to remove any off-axis beams and increases the magnification by a factor 2. Laser wavelengths are separated by dichroic mirrors DC5, DC6 and DC7. Corresponding emission filters C4, C5, C6, and C7 are placed in the emission pathway to cut off any residual excitation light. Additionally, a SH05 Ø1/2" Beam Shutter (Thorlabs) is placed after each emission filter to prevent bleed-through. The wavelengths 485nm and 637nm are separated from 552nm and 738nm and then focused separately on two EMCCD cameras (Xion Ultra 897 EX2 dua AR coated with fringe suppression, Andor Technology Ltd.) to facilitate multiple FRET-pair imaging. Finally, each beam is focused on to the full chip of the EMCCD detectors with aspheric lenses (Qioptiq) with 50.8 mm diameter. Sample chambers designed to integrate temperature, CO2 and humidity control (Okolab, Naples, Italy) are mounted on the microscope stage. For live-cell smFRET imaging, cells were grown in ibidi $\mu$-dish 500 grid glass-bottom plates (ibidi GmbH, Munich). Filters: C1 - FL457.9/10 (ThorLabs), C2 - ZET635/10, C3 - 720/24 BrightLine HC, C4 - 525/50 BrightLine HC, C5 - 675/50 ET, C6 - 593/46 BrightLine HC, C7 - 810/90 ET Bandpass (rest from AHF Analysentechnik). Dichroics: DC1 - BS zt 670 rdc-xxrxt, DC2 - BS zt 561 RDC xrxt, DC3 - BS zt 473 RDC, DC4 - zt488/543/635/730rpc flat, DC5 - T 556 LPXR, DC6 - HC BS 649, DC7 - HC BS 740 (all from AHF Analysentechnik). Notch filters: 458 nm - ZET457NF,638 nm - ZET635NF, 727nm - ZET730NF (all from AHF Analysentechnik).
